## Supplemental Tables and Figures for "Gene regulation modules in non-alcoholic fatty liver disease revealed by single-nucleus ATAC-seq"

**Table S1. Comparison of rat phenotypes between dietary conditions**

| Condition | HFD 4 weeks | HFD 8 weeks | Washout | Normal diet (1) | Normal diet (2) |
|---|---|---|---|---|---|
| Age [weeks] | 16 | 20 | 20 | 16 | 20 |
| Body weight [g] | 292.5 ±13.0 ** | 308.3 ±16.6 ** | 345.2 ±12.2 | 320.8 ±15.4 | 341.2 ±15.2 |
| Liver weight [g] | 15.47 ±0.73 ** | 28.26 ±2.41 ** | 15.33 ±0.59 ** | 10.43 ±0.61 | 11.61 ±0.58 |
| Blood pressure [mmHg] | 171 ±10 ** | 173 ±15 ** | 189 ±14 | 185 ±9 | 191 ±11 |
| Biochemical measurments of serum | | | | | |
| Fasting plasma glucose [mg/dL] | 128.7 ±15.4 ** | 140.6 ±9.1 * | 141.6 ±22.7 | 152.9 ±24.3 | 132.2 ±10.0 |
| Free fatty acids [mEq/l] | 1.33 ±0.29 | 1.08 ±0.32 | 1.57 ±0.33 | 1.14 ±0.26 | 1.30 ±0.25 |
| Triglyceride [mg/dL] | 26.4 ±9.4 | 20.4 ±2.9 | 37.8 ±7.6 | 22.5 ±9.2 | 29.5 ±10.7 |
| Total cholesterol [mg/dL] | 130.9 ±27.0 ** | 120.2 ±19.5 ** | 71.5 ±6.4 * | 74.3 ±6.3 | 79.3 ±3.8 |
| ALT [U/L] | 91.7 ±15.9 ** | 214.6 ±68.5 ** | 45.5 ±7.7 | 42.5 ±3.4 | 47.5 ±3.7 |
| AST [U/L] | 139.2 ±21.0 ** | 421.5 ±102.4 ** | 90.5 ±5.7 | 95.8 ±8.8 | 88.5 ±6.1 |
| $\gamma$-GT [U/L] | 1.60 ±0.51 | 6.88 ±2.29 ** | 1.00 ±0.00 | 1.25 ±0.50 | 1.00 ±0.00 |
| Triglyceride in liver tissue [mg/g] | 21.09 ±4.28 * | 18.87 ±0.97 ** | 15.73 ±4.06 | 13.06 ±1.80 | 11.54 ±1.62 |
| Total cholesterol in liver tissue [mg/g] | 11.32 ±1.63 ** | 17.10 ±0.94 ** | 12.63 ±1.07 ** | 4.17 ±0.98 | 3.29 ±0.34 |
| NAFLD score | | | | | |
| Steatosis [0–3] | 3 [3] | 3 [3] | 1 [1] | 0 [0] | 0 [0] |
| Lobular inflammation [0–3] | 2 [1–2] | 2 [2] | 0 [0] | 0 [0] | 0 [0] |
| Hepatocyte ballooning [0–2] | 1 [0–1] | 1.5 [1–2] | 0 [0] | 0 [0] | 0 [0] |
| NAFLD activity score [0–8] | 6 [4-6] | 6.5 [6–7] | 1 [1] | 0 [0] | 0 [0] |
| Fibrosis stage [0–4] | 0 [0] | 1.5 [1–2] | 0 [0] | 0 [0] | 0 [0] |

Results are shown as Mean ± SD or median [minimum–maximum].

Difference compared to normal diet of same age was tested by t-test: * P<0.05 and ** P<0.01.

Table S2. Prominent genes indicating differential gene expression specific to cell type, induced by dietary intervention. Top three genes are listed. For all genes, see Dataset S1.

| Gene name | Chr | TSS [bp] | HFD 4 weeks Foldchange | HFD 4 weeks P | HFD 8 weeks Foldchange | HFD 8 weeks P | Washout Foldchange | Washout P |
|---|---|---|---|---|---|---|---|---|
| **Hepatocyte** | | | | | | | | |
| Abcg1 | 20 | 9,126,687 | 1.9 | 8.3E-404 | 2.3 | 6.8E-479 | 1.7 | 2.2E-354 |
| Scd | 1 | 243,269,908 | 1.5 | 2.4E-235 | 1.7 | 2.6E-275 | 1.4 | 1.1E-174 |
| Il1r1 | 9 | 42,540,359 | 1.4 | 6.7E-260 | 2.0 | 1.3E-723 | 1.0 | 2.0E-4 |
| Wnt8b | 1 | 243,370,489 | 1.7 | 1.6E-176 | 1.9 | 4.4E-207 | 1.5 | 3.8E-146 |
| Dync2li1 | 6 | 9,992,540 | 2.0 | 4.8E-301 | 2.0 | 2.3E-231 | 1.2 | 1.3E-43 |
| Il1r2 | 9 | 42,384,429 | 1.2 | 3.3E-136 | 1.6 | 3.1E-479 | 1.0 | 8.5E-13 |
| Raph1 | 9 | 61,909,493 | 1.3 | 3.4E-263 | 1.2 | 1.2E-118 | 1.0 | 2.9E-15 |
| LOC259245 | 5 | 75,201,464 | 0.8 | 7.0E-23 | 0.2 | 7.1E-475 | 1.2 | 2.7E-29 |
| Igfbp1 | 14 | 82,047,415 | 0.9 | 3.2E-36 | 1.1 | 2.8E-7 | 0.7 | 1.6E-360 |
| Zfp37 | 5 | 75,598,732 | 0.9 | 1.7E-44 | 0.6 | 1.6E-479 | 1.0 | 1.0E-2 |
| Igfbp3 | 14 | 82,056,347 | 0.7 | 1.1E-114 | 0.9 | 1.0E-3 | 0.4 | 1.2E-644 |
| Bhmt | 2 | 24,859,871 | 0.6 | 6.0E-536 | 0.8 | 1.6E-158 | 0.8 | 8.7E-127 |
| Sqle | 7 | 90,868,742 | 0.7 | 1.2E-403 | 0.8 | 4.6E-261 | 0.8 | 4.0E-430 |
| Mug1 | 4 | 155,076,312 | 0.6 | 4.9E-376 | 0.5 | 1.1E-452 | 0.7 | 1.8E-274 |
| **Endothelial cell** | | | | | | | | |
| Hgf | 4 | 18,677,101 | 1.1 | 5.5E-8 | 1.4 | 4.3E-43 | 1.2 | 9.4E-12 |
| Ackr3 | 9 | 90,799,717 | 1.1 | 3.3E-5 | 1.3 | 1.4E-38 | 1.1 | 2.2E-10 |
| Cfap126 | 13 | 83,526,657 | 1.1 | 4.1E-5 | 1.2 | 4.6E-32 | 1.1 | 1.3E-5 |
| Cemip2 | 1 | 219,345,893 | 1.2 | 3.7E-30 | 1.1 | 3.0E-11 | 1.0 | 6.3E-2 |
| Stab2 | 7 | 21,249,828 | 1.2 | 4.5E-29 | 1.0 | 2.9E-1 | 1.0 | 1.0E-2 |
| Ankrd55 | 2 | 43,914,510 | 1.2 | 3.6E-28 | 1.0 | 1.2E-2 | 1.0 | 3.8E-2 |
| Ice2 | 8 | 70,044,570 | 0.9 | 8.9E-12 | 1.0 | 1.6E-1 | 1.2 | 7.8E-28 |
| Cryba1 | 10 | 62,608,381 | 1.0 | 4.4E-4 | 0.9 | 5.6E-16 | 1.1 | 6.9E-27 |
| Olr1218 | 8 | 38,302,857 | 0.8 | 3.4E-18 | 0.9 | 1.4E-9 | 1.2 | 1.4E-28 |
| Gip | 10 | 80,968,410 | 1.6 | 2.8E-11 | 1.2 | 2.0E-3 | 0.6 | 3.1E-15 |
| Tns3 | 14 | 83,173,649 | 1.1 | 1.5E-6 | 1.0 | 5.6E-5 | 0.9 | 1.3E-23 |
| Il15 | 19 | 25,694,831 | 0.8 | 1.1E-27 | 0.9 | 2.4E-16 | 1.1 | 6.3E-15 |
| Ccr5 | 8 | 123,752,423 | 0.8 | 1.3E-23 | 0.8 | 9.9E-22 | 1.2 | 8.7E-15 |
| Aasdhppt | 8 | 1,453,226 | 0.8 | 6.6E-30 | 0.9 | 3.0E-9 | 1.1 | 1.5E-6 |
| Myo1c | 10 | 60,503,659 | 1.0 | 9.2E-1 | 0.8 | 5.4E-20 | 0.9 | 1.9E-5 |
| Cggbp1 | 11 | 2,565,732 | 0.8 | 1.2E-29 | 0.9 | 1.5E-9 | 1.1 | 2.0E-4 |
| Arc | 7 | 106,555,973 | 0.9 | 3.1E-4 | 0.8 | 4.0E-27 | 1.0 | 1.5E-1 |
| **Stellate cell** | | | | | | | | |
| Epn2 | 10 | 46,197,785 | 1.3 | 5.3E-18 | 1.1 | 4.0E-2 | 1.0 | 1.4E-1 |
| Krt23 | 10 | 84,434,675 | 1.6 | 4.8E-7 | 1.9 | 1.2E-8 | 1.0 | 7.8E-1 |
| Fam71a | 13 | 102,742,988 | 1.6 | 1.0E-12 | 1.3 | 1.1E-2 | 1.0 | 5.4E-1 |
| Scd | 1 | 243,269,908 | 1.1 | 1.4E-2 | 1.3 | 1.5E-8 | 1.1 | 1.0E-1 |
| Adarb1 | 20 | 11,222,595 | 1.1 | 5.8E-3 | 1.2 | 8.6E-9 | 1.0 | 9.3E-1 |
| Pecam1 | 10 | 91,590,521 | 1.1 | 5.4E-15 | 1.0 | 3.9E-2 | 1.0 | 1.3E-2 |
| Faslg | 13 | 74,154,954 | 0.9 | 4.1E-2 | 1.0 | 4.1E-1 | 1.2 | 7.4E-9 |
| Gli3 | 17 | 49,438,567 | 1.0 | 3.9E-1 | 0.9 | 1.2E-1 | 1.3 | 1.1E-9 |
| Cyp2j3 | 5 | 111,220,046 | 1.0 | 2.4E-1 | 0.8 | 4.7E-4 | 1.3 | 1.7E-10 |
| Amotl2 | 8 | 103,303,755 | 1.0 | 2.3E-1 | 0.8 | 1.3E-10 | 1.0 | 9.4E-2 |
| Fbln5 | 6 | 120,899,220 | 0.9 | 1.3E-3 | 0.8 | 4.5E-10 | 1.1 | 3.8E-5 |
| Msmo1 | 16 | 24,980,680 | 0.8 | 6.1E-17 | 1.0 | 7.5E-1 | 1.1 | 1.6E-2 |
| Rgr | 16 | 12,801,594 | 0.8 | 6.0E-15 | 0.9 | 1.2E-1 | 1.1 | 4.4E-3 |
| Nlrp4 | 1 | 68,089,321 | 0.9 | 2.7E-1 | 0.9 | 1.6E-1 | 0.7 | 8.0E-7 |
| Mup4 | 5 | 74,961,236 | 0.9 | 6.0E-1 | 0.8 | 7.5E-2 | 0.6 | 2.8E-7 |
| RGD1305938 | 2 | 53,189,789 | 0.8 | 1.8E-14 | 0.9 | 4.1E-2 | 1.0 | 8.3E-1 |
| Zfp37 | 5 | 75,598,732 | 0.9 | 2.8E-4 | 0.8 | 3.8E-10 | 1.0 | 4.2E-1 |
| Cyp2c6v1 | 1 | 237,693,095 | 0.8 | 3.2E-5 | 0.8 | 5.1E-3 | 0.8 | 5.4E-8 |
| **Macrophage** | | | | | | | | |
| RT1-Db2 | 20 | 4,521,048 | 2.5 | 1.6E-36 | 2.1 | 2.9E-25 | 2.3 | 5.2E-23 |
| RT1-Da | 20 | 4,513,485 | 2.2 | 2.6E-42 | 1.7 | 5.0E-22 | 1.8 | 2.3E-20 |
| Scd | 1 | 243,269,908 | 1.6 | 5.5E-47 | 1.5 | 1.1E-47 | 1.1 | 5.6E-3 |
| RT1-Db1 | 20 | 4,548,664 | 2.1 | 1.6E-41 | 1.5 | 1.5E-12 | 1.5 | 2.4E-10 |
| Fabp9 | 2 | 91,592,306 | 2.2 | 5.5E-20 | 3.4 | 6.5E-40 | 0.9 | 9.8E-2 |
| LOC102550325 | 2 | 174,693,445 | 1.1 | 5.9E-3 | 1.7 | 4.2E-40 | 1.1 | 3.2E-3 |
| Klra2 | 4 | 164,950,581 | 1.1 | 2.3E-2 | 1.1 | 2.9E-1 | 1.6 | 1.4E-19 |
| Tns1 | 9 | 75,495,801 | 1.1 | 5.2E-8 | 1.0 | 3.3E-4 | 0.8 | 1.1E-22 |
| Cdc42ep4 | 10 | 98,772,613 | 1.0 | 1.3E-1 | 1.0 | 3.6E-3 | 0.8 | 9.9E-17 |
| Npepo | 17 | 1,815,405 | 1.0 | 1.2E-1 | 1.0 | 6.3E-2 | 0.9 | 2.3E-16 |
| Rasgrp2 | 1 | 203,707,481 | 0.9 | 1.2E-3 | 0.7 | 2.2E-23 | 0.9 | 2.9E-2 |
| Arhgap5 | 6 | 69,988,210 | 0.8 | 5.5E-44 | 0.9 | 4.2E-4 | 1.0 | 9.3E-1 |
| Crybb3 | 12 | 43,557,103 | 0.8 | 1.6E-22 | 0.8 | 8.7E-23 | 1.0 | 3.3E-1 |
| Gsta6 | 9 | 23,743,262 | 0.7 | 7.5E-46 | 0.8 | 4.1E-20 | 1.0 | 4.6E-1 |
| Ehd3 | 6 | 21,640,861 | 0.8 | 2.8E-18 | 0.8 | 4.4E-24 | 0.9 | 1.1E-5 |
| Fbxl21 | 17 | 8,061,292 | 0.6 | 7.5E-50 | 0.7 | 3.4E-17 | 0.9 | 9.5E-2 |

**Table S3. Potential drug repositioning for NAFLD**

| gene_name | interaction_claim_source | interaction_types | drug_claim_name | drug_claim_primary_name | drug_name | drug_concept_id | interaction_group_score |
|---|---|---|---|---|---|---|---|
| **Antagonist/antibody/inhibitor of upregulated or positively correlated genes of NAFLD** | | | | | | | |
| BCL2L1 | GuideToPharmacology | antagonist | 252166532 | ABT-737 | ABT 737 | chembl:CHEMBL376408 | 4.55 |
| BCL2L1 | GuideToPharmacology | antagonist | 252166530 | VENETOCLAX | VENETOCLAX | chembl:CHEMBL3137309 | 0.57 |
| BCL2L1 | GuideToPharmacology | antagonist | 252166531 | NAVITOCLAX | NAVITOCLAX | chembl:CHEMBL443684 | 2.65 |
| BCL2L1 | ChemblInteractions | inhibitor | CHEMBL443684 | NAVITOCLAX | NAVITOCLAX | chembl:CHEMBL443684 | 2.65 |
| BCL2L1 | TALC | inhibitor | NAVITOCLAX | NAVITOCLAX | NAVITOCLAX | chembl:CHEMBL443684 | 2.65 |
| BCL2L1 | ChemblInteractions | inhibitor | CHEMBL2107358 | OBATOCLAX MESYLATE | OBATOCLAX MESYLATE | chembl:CHEMBL2107358 | 1.33 |
| CXCR4 | TALC | antagonist | LY 2510924 | LY 2510924 | | | NA |
| CXCR4 | ChemblInteractions | antagonist | CHEMBL3545330 | MSX-122 | MSX-122 | chembl:CHEMBL3545330 | 9.55 |
| CXCR4 | ChemblInteractions | antagonist | CHEMBL3545105 | POL6326 | POL6326 | chembl:CHEMBL3545105 | 6.37 |
| CXCR4 | GuideToPharmacology | antagonist | 252166781 | MAVORIXAFOR | MAVORIXAFOR | chembl:CHEMBL518924 | 15.91 |
| CXCR4 | GuideToPharmacology | antagonist | 135650433 | ISOTHIOUREA-1T | | | NA |
| CXCR4 | GuideToPharmacology | antagonist | 135652593 | T140 | BKT140 | chembl:CHEMBL3545348 | 12.73 |
| CXCR4 | GuideToPharmacology | antagonist | 363894176 | TIQ-15 | | | NA |
| CXCR4 | GuideToPharmacology | antagonist | 135652455 | SDF-1, 1-9[P2G] DIMER | | | NA |
| CXCR4 | GuideToPharmacology | antagonist | 135649932 | PLERIXAFOR | PLERIXAFOR | chembl:CHEMBL18442 | 8.91 |
| CXCR4 | GuideToPharmacology | antagonist | 135652187 | HIV-TAT | | | NA |
| CXCR4 | GuideToPharmacology | antagonist | 404859126 | MOTIXAFORTIDE | BKT140 | chembl:CHEMBL3545348 | 12.73 |
| CXCR4 | ChemblInteractions | antagonist | CHEMBL3544949 | CTCE-9908 | CTCE-9908 | chembl:CHEMBL3544949 | 6.37 |
| CXCR4 | GuideToPharmacology | antagonist | 135652592 | T134 | | | NA |
| CXCR4 | GuideToPharmacology | antagonist | 135650432 | ISOTHIOUREA-1A | | | NA |
| CXCR4 | ChemblInteractions | antagonist | CHEMBL3545224 | Burixafor | BURIXAFOR | chembl:CHEMBL3545224 | 9.55 |
| CXCR4 | GuideToPharmacology | antagonist | 135652636 | VIRAL MACROPHAGE INFLAMMATORY PROTEIN-II | | | NA |
| CXCR4 | ChemblInteractions | antagonist | CHEMBL3545348 | BKT140 | BKT140 | chembl:CHEMBL3545348 | 12.73 |
| CXCR4 | GuideToPharmacology | antagonist | 348353660 | CX549 | | | NA |
| CXCR4 | GuideToPharmacology | antagonist | 381118856 | CXCR4 ANTAGONIST 22 | | | NA |
| CXCR4 | GuideToPharmacology | antagonist | 363894177 | COMPOUND 46C [PMID: 29350534] | | | NA |
| CXCR4 | GuideToPharmacology | antagonist | 135652594 | T22 | | | NA |
| CXCR4 | GuideToPharmacology | antagonist | 252166738 | SDF1 P2G | | | NA |
| CXCR4 | GuideToPharmacology | antibody | 315661244 | ULOCUPLUMAB | ULOCUPLUMAB | chembl:CHEMBL3039543 | 6.37 |
| HGF | TALC | antibody | RILOTUMUMAB | RILOTUMUMAB | RILOTUMUMAB | chembl:CHEMBL1743063 | 13.26 |
| HGF | TALC | antibody | FICLATUZUMAB | FICLATUZUMAB | FICLATUZUMAB | chembl:CHEMBL1743018 | 13.26 |
| HGF | MyCancerGenome | antibody | FICLATUZUMAB | FICLATUZUMAB | FICLATUZUMAB | chembl:CHEMBL1743018 | 13.26 |
| HGF | MyCancerGenome | antibody | RILOTUMUMAB | RILOTUMUMAB | RILOTUMUMAB | chembl:CHEMBL1743063 | 13.26 |
| HGF | ChemblInteractions | inhibitor | CHEMBL1743063 | RILOTUMUMAB | RILOTUMUMAB | chembl:CHEMBL1743063 | 13.26 |
| HGF | CancerCommons | inhibitor | AMG 102 | AMG 102 | RILOTUMUMAB | chembl:CHEMBL1743063 | 13.26 |
| HGF | ChemblInteractions | inhibitor | CHEMBL1743018 | FICLATUZUMAB | FICLATUZUMAB | chembl:CHEMBL1743018 | 13.26 |
| IL1R1 | ChemblInteractions | antagonist | CHEMBL1201570 | ANAKINRA | ANAKINRA | chembl:CHEMBL1201570 | 17.24 |
| IL1R1 | ChemblInteractions | antagonist | CHEMBL2109458 | AMG-108 | AMG-108 | chembl:CHEMBL2109458 | 5.3 |
| TNF | ChemblInteractions | inhibitor | CHEMBL2107911 | ONERCEPT | ONERCEPT | chembl:CHEMBL2107911 | 0.94 |
| TNF | ChemblInteractions | inhibitor | CHEMBL2109587 | AZ-9773 | AZ9773 | chembl:CHEMBL2109587 | 0.94 |
| TNF | ChemblInteractions | inhibitor | CHEMBL1201831 | CERTOLIZUMAB PEGOL | CERTOLIZUMAB PEGOL | chembl:CHEMBL1201831 | 0.94 |
| TNF | ChemblInteractions | inhibitor | CHEMBL2108887 | AFELIMOMAB | AFELIMOMAB | chembl:CHEMBL2108887 | 1.87 |
| TNF | ChemblInteractions | inhibitor | CHEMBL1201580 | ADALIMUMAB | ADALIMUMAB | chembl:CHEMBL1201580 | 1.43 |
| TNF | ChemblInteractions | inhibitor | CHEMBL2108566 | NERELIMOMAB | NERELIMOMAB | chembl:CHEMBL2108566 | 0.94 |
| TNF | ChemblInteractions | inhibitor | CHEMBL2108208 | LENERCEPT | LENERCEPT | chembl:CHEMBL2108208 | 0.94 |
| TNF | ChemblInteractions | inhibitor | CHEMBL1743054 | OZORALIZUMAB | OZORALIZUMAB | chembl:CHEMBL1743054 | 1.87 |
| TNF | ChemblInteractions | inhibitor | CHEMBL2108739 | PLACULUMAB | PLACULUMAB | chembl:CHEMBL2108739 | 2.81 |
| TNF | ChemblInteractions | inhibitor | CHEMBL1743057 | PEGSUNERCEPT | PEGSUNERCEPT | chembl:CHEMBL1743057 | 1.87 |
| TNF | ChemblInteractions | inhibitor | CHEMBL1201581 | INFLIXIMAB | INFLIXIMAB | chembl:CHEMBL1201581 | 1.59 |
| TNF | ChemblInteractions | inhibitor | CHEMBL1201833 | GOLIMUMAB | GOLIMUMAB | chembl:CHEMBL1201833 | 4.68 |
| TNF | ChemblInteractions | inhibitor | CHEMBL1201572 | ETANERCEPT | ETANERCEPT | chembl:CHEMBL1201572 | 1.28 |
| **Agonist of downregulated or negatively correlated genes of NAFLD** | | | | | | | |
| CCR5 | GuideToPharmacology | agonist | 135651687 | CCL4 | | | NA |
| CCR5 | GuideToPharmacology | agonist | 178100743 | R5-HIV-1 GP120 | | | NA |
| CCR5 | GuideToPharmacology | agonist | 135651657 | CCL13 | | | NA |
| CCR5 | GuideToPharmacology | agonist | 135651654 | CCL11 | | | NA |
| CCR5 | GuideToPharmacology | agonist | 135652155 | FLU-CCL3 | | | NA |
| CCR5 | GuideToPharmacology | agonist | 135651684 | CCL3 | | | NA |
| CCR5 | GuideToPharmacology | agonist | 135651690 | CCL5 | | | NA |
| CCR5 | GuideToPharmacology | agonist | 135652497 | [125I]CCL4 (HUMAN) | | | NA |
| CCR5 | GuideToPharmacology | agonist | 135651660 | CCL16 | | | NA |
| CCR5 | GuideToPharmacology | agonist | 135652498 | [125I]CCL5 (HUMAN) | | | NA |
| CCR5 | GuideToPharmacology | agonist | 135652496 | [125I]CCL3 (HUMAN) | | | NA |
| CCR5 | GuideToPharmacology | agonist | 135651694 | CCL8 | | | NA |
| CCR5 | GuideToPharmacology | agonist | 135652500 | [125I]CCL8 (HUMAN) | | | NA |
| CCR5 | GuideToPharmacology | agonist | 135651664 | CCL2 | | | NA |
| CCR5 | GuideToPharmacology | agonist | 135651658 | CCL14 | | | NA |
| CCR5 | GuideToPharmacology | agonist | 135652047 | BP-CCL3 | | | NA |
| CXCR4 | GuideToPharmacology | agonist | 252166735 | CXCL12Φ | | | NA |
| CXCR4 | GuideToPharmacology | agonist | 135652070 | CXCL12-(1-9) | | | NA |
| CXCR4 | GuideToPharmacology | agonist | 135651882 | CXCL12B | | | NA |
| CXCR4 | GuideToPharmacology | agonist | 135652069 | CXCL12-(1-17) | | | NA |
| CXCR4 | GuideToPharmacology | agonist | 178100751 | X4-HIV-1 GP120 | | | NA |
| CXCR4 | GuideToPharmacology | agonist | 135651987 | ALX40-4C | | | NA |
| CXCR4 | GuideToPharmacology | agonist | 135652071 | CXCL12-(1-9) DIMER | | | NA |
| CXCR4 | GuideToPharmacology | agonist | 252166736 | CXCL12H25R (MONOMER) | | | NA |
| CXCR4 | GuideToPharmacology | agonist | 135652555 | [125I]CXCL12A (HUMAN) | | | NA |
| CXCR4 | GuideToPharmacology | agonist | 252166733 | CXCL12Δ | | | NA |
| CXCR4 | GuideToPharmacology | agonist | 135652556 | [125I]CXCL12B (HUMAN) | | | NA |
| CXCR4 | GuideToPharmacology | agonist | 252166734 | CXCL12E | | | NA |
| CXCR4 | GuideToPharmacology | agonist | 252166737 | CXCL122 (DIMER) | | | NA |
| CXCR4 | GuideToPharmacology | agonist | 252166732 | CXCL12Γ | | | NA |
| CXCR4 | GuideToPharmacology | agonist | 135651881 | CXCL12A | | | NA |
| TLR2 | GuideToPharmacology | agonist | 381745022 | DIPROVOCIM-1 | | | NA |
| TLR2 | GuideToPharmacology | agonist | 310264669 | COMPOUND 13 [PMID: 23098072] | | | NA |
| TLR2 | GuideToPharmacology | agonist | 178101739 | PEPTIDOGLYCAN | | | NA |
| TLR4 | GuideToPharmacology | agonist | 381118801 | CRX-555 | | | NA |
| TLR4 | GuideToPharmacology | agonist | 178101716 | LPS | | | NA |
| TLR4 | GuideToPharmacology | agonist | 315661222 | NEOCEPTIN-3 | | | NA |

Figure S1

**Cross-check of differential gene expression (DGE) between single-nucleus assay in this study and microarray experiment of bulk tissue**

DGE was computed by comparing the 4 weeks of HFD condition with the normal diet condition. The horizontal axis indicates the z-score for cell type-specific DGE observed in this study using snATAC-seq. The vertical axis indicates the z-score for DGE observed in the Agilent Rat GE 4x44K v3 Microarray experiment of bulk liver tissue samples (n=15 and 12 for 4 weeks of HFD and normal diet, respectively). The blue line shows the smoothed fit, and the gray area shows the 95% confidence interval.

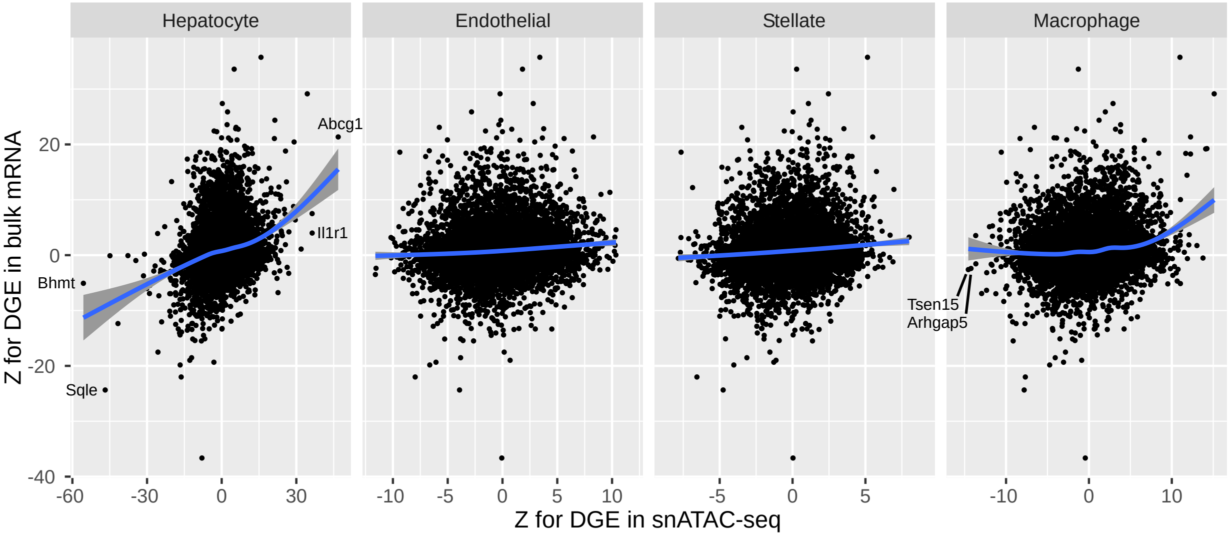

Figure S2

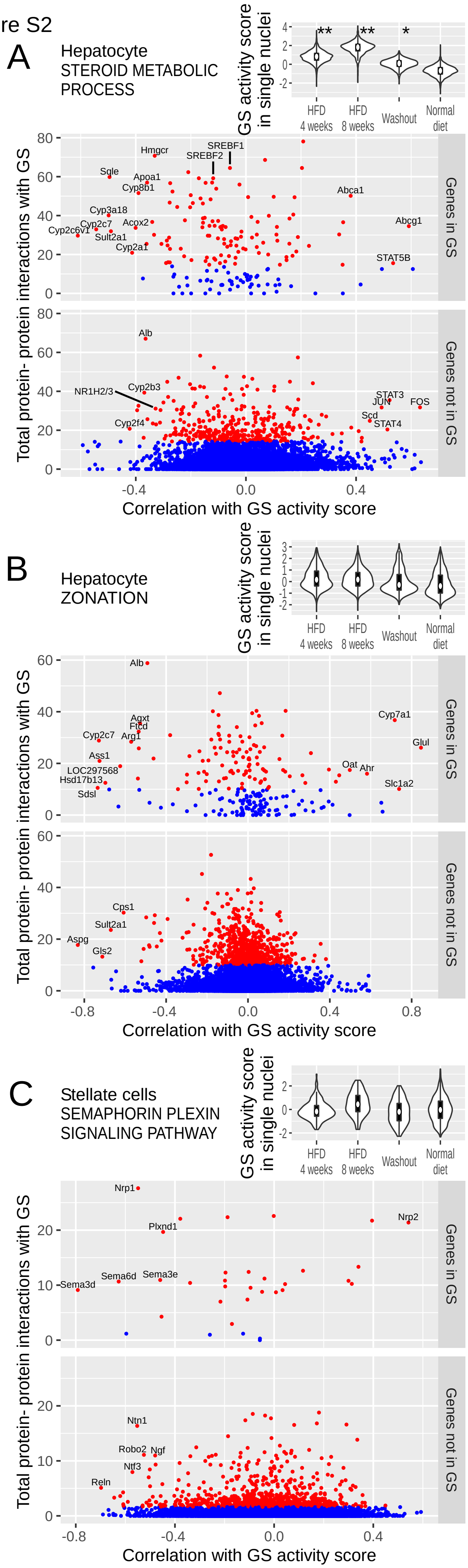


**Core genes for biological processes other than inflammation are extracted based on correlation with GS activity score and protein-protein interaction**

(A) Analysis of hepatocytes using the GS for steroid metabolism, taken from Gene Ontology database. See the legend for Figure 7A.

(B) Analysis of hepatocytes using the GS for zonation in liver lobule.

(C) Analysis of stellate cells using the GS for semaphorin-plexin signaling, taken from the Gene Ontology database.

Figure S3

**The effect of machine learning hyperparameters on TF module discovery**
TF modules were computed in two steps, first the regulator-regulatee matrix was computed using GENIE3, then the modules were extracted by nonnegative matrix factorization using NMF. We assessed the accuracy of TF module discovery under a range of hyperparameters by randomly dividing the hepatocytes into two sets of halves (4136 nuclei in each set), computing in each set the regulator-regulatee matrix, approximating the matrix using NMF (with nonnegative double singular value decomposition seed), and then calculating the correlation coefficient between the two NMF-approximated regulator-regulatee matrices. A larger correlation coefficient indicates a higher accuracy. To estimate the noise level, we performed the same process in each set but with nuclei labeling of the Gene Score Matrix randomly permuted while keeping the Motif Matrix intact. Under each setting of the hyperparameter, we performed five permutation trials. The correlation coefficient is plotted as dots and its mean plus/minus standard deviation is plotted as boxes.
(A) As for the number of candidate regulators, *K*, randomly selected at each node of the random forests, we examined 1, 23 (the adopted value, which is the square root of the number of regulators) and 30. The correlation coefficient gradually decreased as *K* increased, but was close to one and much larger than the permuted trials.
(B) As for the rank of NMF, we examined 2, 4 (the adopted value), 6 and 8. The correlation coefficient was similarly high for 2 and 4, but dropped from 6, indicating overfitting in the latter.

A

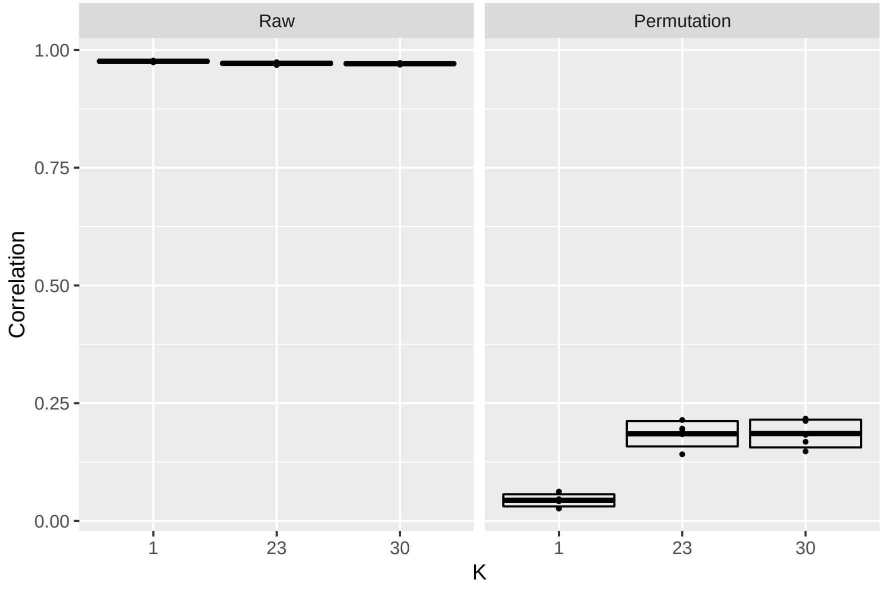


B

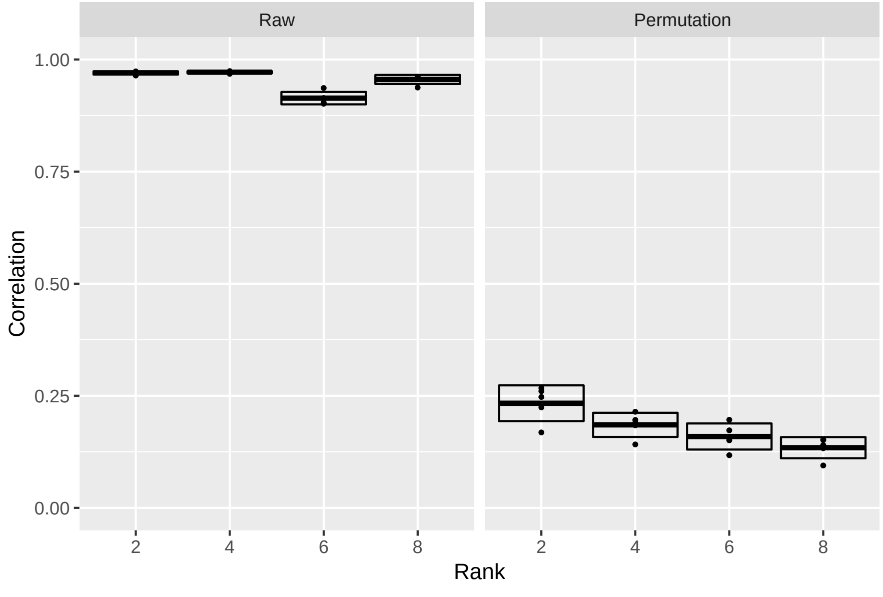

Figure S4

**The effect of GRN algorithms on TF module discovery**
As GRN identification algorithms vary in their definitions of GRN (Figure 9), we compared SCENIC+ and CellOracle with our algorithm. Our algorithm ’ s TF-gene linking criterion is relatively relaxed, while SCENIC+ applies a more stringent criterion. Our algorithm requires correlation between ATAC-seq peak accessibility and gene expression, which CellOracle does not require. First, the GRN of hepatocytes was computed using each respective algorithm. Then, the same module discovery method was applied to each GRN: four major components were extracted via nonnegative matrix factorization, and relevant biological processes were detected by gene set enrichment analysis. In SCENIC+, module 3 displayed a strong signal for steroid metabolism, while the other modules presented modest signals for cell adherence, innate immunity and inflammation. CellOracle did not reveal any statistically significant biological process. Our algorithm identified three modules with strong signals for steroid metabolism, inflammation, and lobular zonation.

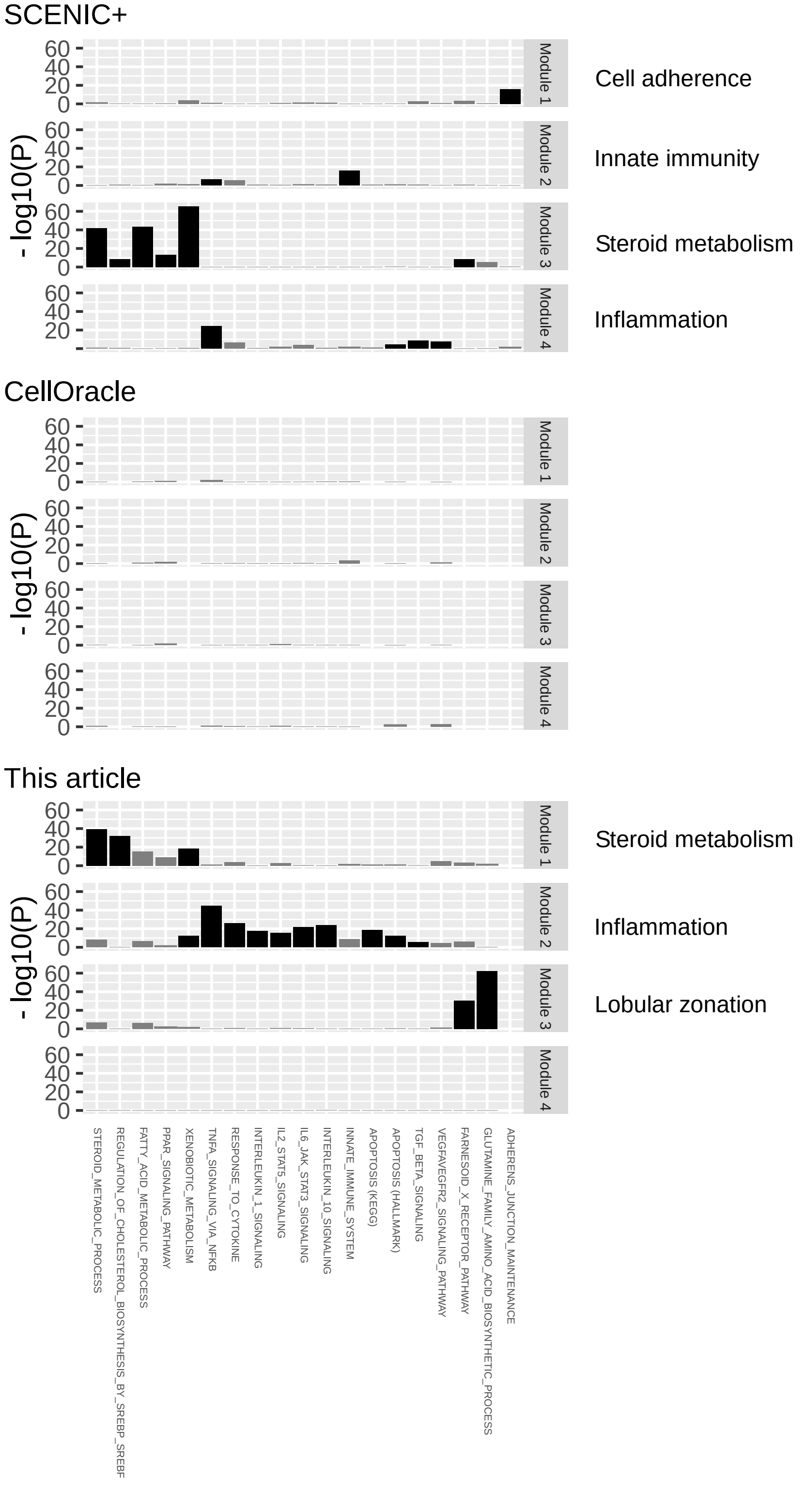
